# Optimizing Genome Editing in Bovine Cells: A Comparative Study of Cas9 Variants and CRISPR Delivery Methods

**DOI:** 10.1101/2024.11.07.620146

**Authors:** Xiaofeng Du, Alexander Quinn, Timothy Mahony, Di Xia, Laercio R. Porto-Neto

## Abstract

To assist in the establishment of an efficient CRISPR/Cas9 gene editing workflow in bovine cells, we compared the efficiency of four *Streptococcus pyogenes* Cas9 (SpCas9) nuclease variants (produced in-house or commercially) and two different Cas9/sgRNA ribonucleoprotein (RNP) delivery methods applied to Madin-Darby bovine kidney (MDBK) cells (*Bos taurus*). We targeted three genes for simple sequence deletion or modification via a single-stranded oligodeoxynucleotide donor template: the testis-determining gene *Sry* (sex-determining region on Y chromosome), germ cell-specific gene *Nanos2* (nanos C2HC-type zinc finger 2) and *PRLR* (prolactin receptor). RNPs and donor templates were delivered into cells via lipofectamine CRISPRMAX transfection or Neon electroporation. The efficiency of gene editing was determined by target-specific PCR genotyping, real-time PCR assays and next generation sequencing analyses. When targeting *Sry*, the commercial Alt-R High-Fidelity (HiFi) SpCas9 nuclease exhibited the highest deletion efficiency, followed by the in-house generated Sniper2L, HiFi SpCas9 and wild-type SpCas9. Notably, for *PRLR* and *Nanos2*, Sniper2L induced comparable editing outcomes to Alt-R HiFi SpCas9. The two delivery methods, lipofectamine CRISPRMAX transfection and Neon electroporation, demonstrated similar efficiency (60% to 83%) in producing indels in all three target genes. However, Neon electroporation (5.5% to 11%) was superior to CRISPRMAX lipofection (1.5% to 4.8%) at inducing target-specific sequence knock-ins. Strategies to inhibit NHEJ repair and/or enhance HDR may be necessary to improve HDR efficiency in these bovine cells. These findings provide valuable insights for improving gene editing outcomes in bovines and may assist in accelerating the widespread application of genome editing technology in large animals.

## 1. Introduction

Genome editing has been applied to livestock for diverse purposes, including improving the efficiency and environmental sustainability of food production, enhancing animal welfare and disease resistance, and generating animal models for biomedical research [1–3]. The CRISPR (Clustered Regularly Interspaced Short Palindromic Repeats)/Cas9 (CRISPR-associated protein 9) system is a Nobel Prize-winning molecular technique that has revolutionized the field of gene editing. The versatility, high efficiency and specificity of CRISPR/Cas9 provide considerable advantages over other genome engineering tools, such as engineering zinc finger nucleases and transcription activator-like effector nucleases [4–9]. In the CRISPR/Cas9 gene editing system, the Cas9 ribonuclease is complexed with a single guide RNA (sgRNA) that includes a short (∼20 nucleotide) sequence complementary to a desired target DNA sequence. Once the Cas9-sgRNA ribonucleoprotein (RNP) complex binds to the target DNA site, the Cas9 nuclease cleaves the DNA, introducing a site-specific double strand break (DSB) [10,11]. In cells, DSBs are typically repaired through a process of non-homologous end joining (NHEJ) which randomly introduces small insertions or deletions (indels) of nucleotides at the repaired site. An alternative mechanism of homology-directed repair (HDR) can also occur (albeit at lower frequency than NHEJ) if a donor DNA template with sequence homologous to both sides of the DSB is available at the time of repair, resulting in highly precise DNA repair of the DSB or even introduction (‘knock-in’) of a desired novel sequence at the targeted site [12–14].

To date, two main strategies have been applied to generate gene-edited large animals: (a) direct genome editing of zygotes [15–18]; and (b) editing of somatic cells, which are then used to generate an embryo via somatic cell nuclear transfer (SCNT) [17,19–22]. Each approach has advantages and disadvantages [23–25]. Direct editing of zygotes via cytoplasmic microinjection or electroporation of gene editing components has minimal impact on embryo development and is more efficient at producing viable animals [17,25–27]. However, it is difficult to control and characterize gene editing events in embryos following the introducing of genome editors such as Cas9. Consequently, edited embryos often have a mosaic pattern of edited cells or exhibit unintended off-target sequence edits [1,17,26,28]. In contrast, somatic cell-mediated gene editing has the advantage of allowing full characterization of gene edits, and selection of correctly edited cells for the generation of embryos through SCNT. However, the production of live animals from SCNT remains very inefficient, and live-born animals may have reduced viability [22,23,29,30]. Notwithstanding the application, efficient gene editing workflows are required to shorten the time it takes to obtain viable animals with the desired genotype, to reduce costs of this process, and to improve animal welfare. This is especially true for the application of gene editing approaches in large animals like cattle which have a relatively long generation interval and low fecundity.

Herein, as part of an effort to establish an efficient CRISPR/Cas9 gene editing workflow for bovines, we compared the efficiency of four different Cas9 nuclease variants and two Cas9 RNP delivery methods (lipofection and electroporation) for inducing targeted gene knockout and knock-in in Madin-Darby bovine kidney (MDBK) cells [31]. We targeted three key genes in *Bos taurus*: *Sry* (sex-determining region on Y chromosome), germ cell-specific gene *Nanos2* (nanos C2HC-type zinc finger 2) and *PRLR* (prolactin receptor). Gene editing outcomes were analysed by endpoint and real-time PCR assays, Sanger sequencing, and next generation sequencing (NGS).

## 2. Methods and materials

### 2.1. Design of sgRNAs and HDR donors

SgRNAs were designed using the online design tool Benchling (https://benchling.com/) [32]. The sgRNA sequences were also analyzed in BLAST to confirm their specificity. Single-stranded oligonucleotide HDR donor templates were designed with the assistance of Edit-R HDR Donor Designer-oligo (Horizon Discovery, United Kingdom). For editing of *Sry*, two sgRNAs (*Sry*-sgRNA1 and *Sry*-sgRNA2) were selected to delete the high-mobility group (HMG) domain (350 bp) (**Figure 1A**, **Supplementary Table S1**). For knock-in at the *Sry* locus, we designed a 104 nt single-stranded oligodeoxynucleotide (ssODN) donor template, with homology arms of 40 nt flanking a central 24 nt sequence encoding six stop codons (two for each possible reading frame; 5’-<u>TAA</u>G<u>TGA</u>C<u>TAG</u>G<u>TAA</u>C<u>TGA</u>G<u>TAG</u>C-3’) (**Figure 1A**, **Supplementary Table S1**). The homology arms of the ssODN donor were designed to insert the 24 nt transgenic sequence into the DSB induced by *Sry-*sgRNA1. Additionally, to determine the most effective type of donor template for targeted knock-in to bovine cells, we tested three formats of ssODN donors commercially available from Integrated DNA Technologies: Alt-R HDR modification (donor 1), phosphorothioate (PS) bond modification (donor 2) and non-modification (donor 3).

**Figure 1.**
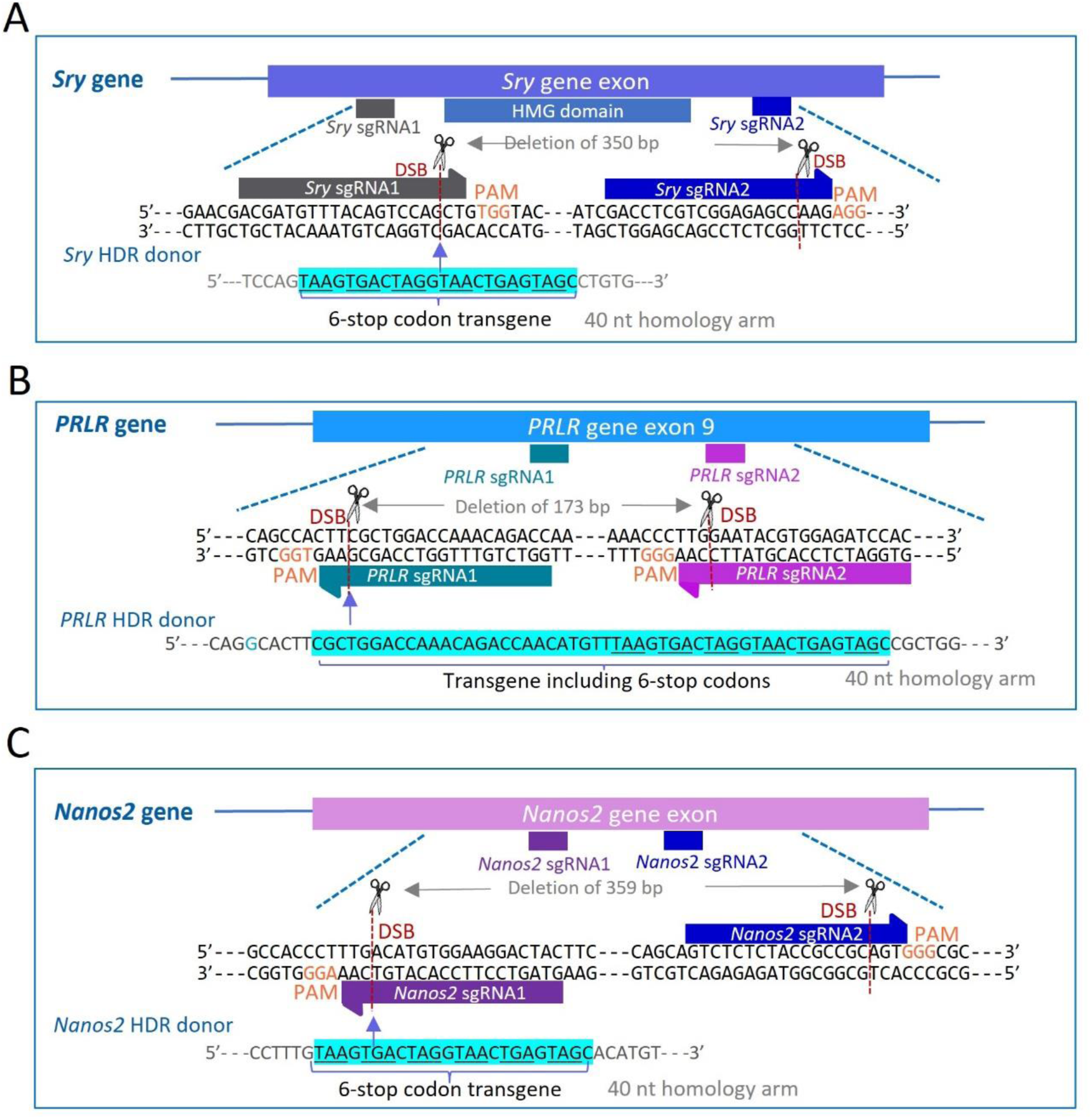
CRISPR/Cas9 gene editing strategy for *Bos taurus* genes *Sry*, *PRLR* and *Nanos2*. Protospacer adjacent motifs (PAMs) for sgRNA binding sites are indicated by light brown text, and predicted sites for Cas9-induced double-strand breaks (DSBs) are indicated by dashed red lines. (**A**) Binding site locations for *Sry* sgRNA1 and *Sry* sgRNA2, flanking the high mobility group (HMG) domain of the *Sry* gene. The *Sry* HDR donor template (24 nt 6-stop codon transgene flanked by 40 nt homology arms) is designed to insert into the DSB generated by *Sry* sgRNA1. (**B**) Binding sites and predicted DSB locations for *PRLR* sgRNA1 and sgRNA2. The *PRLR* HDR donor template (50 nt transgene flanked by 40 nt homology arms) is designed to insert into the DSB generated by *PRLR* sgRNA1. A mutation was introduced into the PAM sequence in the HDR donor was modified at one nucleotide to protect the donor DNA from Cas9 cleavage. (**C**) Binding sites and predicted DSB locations for *Nanos2* sgRNA1 and sgRNA2 [17]. The *Nanos2* HDR donor template (24 nt 6-stop codon transgene flanked by 40 nt homology arms) is designed to insert into the DSB created by *Nanos2* sgRNA1.

*PRLR*-sgRNA1 and *PRLR*-sgRNA2 were designed to delete 173 bp from exon 9 of *PRLR* and to create a stop codon at the target site (**Figure 1B**, **Supplementary Table S1**). A 130 nt *PRLR* specific ssODN (40 nt homology arms flanking a 50 nt transgenic insertion including the same 24 nt sequence encoding six stop codons) was designed as an HDR donor template (Alt-R HDR modification) (**Figure 1B**, **Supplementary Table S1**). The 50 nt transgene was designed to insert into the *PRLR-*sgRNA1 cleavage site and modify the sequence to mimic the natural mutation of SLICK cattle [33,34]. The *PRLR* donor template also replaced the TGG PAM sequence in the genomic sequence with TGC, to protect the HDR donor DNA and the integrated transgenic sequence from Cas9 cleavage. The *Nanos2*-targeting sgRNAs (*Nanos2*-sgRNA1 and *Nanos2*-sgRNA2) we used were previously published by Miao *et al.* [17]. In addition, we designed a *Nanos2*-specific ssODN donor (Alt-R HDR modification) for targeted knock-in of the six-stop-codon transgene into the *Nanos2*-sgRNA1 target locus (**Figure 1C**, **Supplementary Table S1**). The designed sgRNAs and ssODN donors were synthesized by Integrated DNA Technologies (IDT, Coralville, Iowa, USA).

### 2.2. Construction of DNA plasmids for bacterial expression of Cas9 nucleases

The 4.1 kb coding sequences for Cas9 variants wild-type (WT) SpCas9, High-Fidelity (HiFi) SpCas9 and Sniper2L were amplified by HiFi PCR using plasmids #52961, #188490 and #193856 (Addgene, USA), respectively, as templates. The four plasmids were gifts from David Liu (#62933 [35]), Feng Zhang (#52961 [36]), Isaac Hilton (#188490 [37]) and Hyongbum Kim (#193856 [38]). A pET bacterial expression vector (plasmid #62933, Addgene, USA) was digested with restriction enzymes *Pac*I and *Xba*I (New England Biolabs, Ipswich, MA, USA), and the 5.0 kb backbone (including T7 RNA polymerase promoter and terminator sequences) was purified from the digest by gel extraction. The amplified Cas9 sequences were purified and cloned into the purified 5.0 kb pET plasmid backbone, using the NEBuilder® HiFi DNA Assembly Cloning Kit (New England Biolabs). The four-fragment Gibson [39] Assembly reactions included an 89 bp single-stranded oligo designed to bridge the N-terminal end of the Cas9 sequence with the T7 promoter end of the pET backbone, and a 276 bp double-stranded gBlock™ fragment (IDT) designed to bridge the C-terminal end of the Cas9 sequence with the T7 terminator end of the backbone. The gBlock fragment also incorporated sequences for three C-terminal nuclear localization signal (NLS) peptides (1× Nucleoplasmin NLS, 2× c-Myc NLS) a 6× Histidine tag (to facilitate protein purification), and three successive stop codons. The 3× NLS structure was designed to improve gene editing efficacy of the Cas9 variants relative to Cas9 protein with a single NLS [40]. The three newly constructed plasmids thus coded for proteins SpCas9-3×NLS-6×His, HiFi-SpCas9-3×NLS-6×His and Sniper2L-3×NLS-6×His. The plasmids were transformed into *E. coli* DH5α cells (New England Biolabs), and endotoxin-free plasmid DNA was purified from 100 ml of cultured cells using the NucleoBond Xtra Midi Plus EF kit (Macherey Nagel, Germany). The full coding sequence of 1,426 amino acids was verified by Sanger sequencing for all three plasmids.

### 2.3. Expression and purification of Cas9 variant proteins

Alt-R S.p. HiFi SpCas9 3NLS nuclease was purchased from IDT. Recombinant Sniper2L (a new HiFi SpCas9 variant [38]), HiFi SpCas9 [37], and WT SpCas9 [36] were expressed and purified in the lab [41]. Briefly, the engineered vectors for each protein were transformed into *Escherichia coli* BL21 DE3 strain (Sigma-Aldrich, Melbourne, Australia). The transformed *E. coli* bacteria were plated on antibiotic selection (ampicillin^+^) LB agar plates and incubated at 37°C overnight. Subsequently, a single colony was used to inoculate 10 ml LB media with ampicillin (100 μg/ml), which was then incubated overnight at 37°C with shaking at 220 RPM. The bacterial culture was added to 1 L fresh LB media containing 100 μg/ml ampicillin and further cultured (37°C, 220 RPM) until OD_600_ reached 0.4-0.8. IPTG (isopropyl-β-D-thiogalactopyranoside) (Sigma-Aldrich) was added to the bacteria culture to a final concentration of 0.5 mM to induce protein expression at 16°C with shaking at 220 RPM for 20 h. Bacterial cells were pelleted by centrifugation at 4,500 g for 20 min at 4°C.

The harvested bacteria cells were resuspended in 30 ml lysis buffer [50 mM Tris-HCl, 500 mM NaCl, 5 mM MgCl_2_, pH 8.0, 100 μg/ml of lysozyme (Thermo Fisher Scientific, Brisbane, Australia), 25 U/ml of Pierce Universal Nuclease (Thermo Fisher Scientific), and SIGMAFAST™ Protease Inhibitor Cocktail (Sigma-Alrich)]. The cell suspension was then lysed using a Misonix Microson Ultrasonic Cell Disruptor (Farmingdale, New York, United States), followed by centrifugation at 16 000 g for 30 min at 4°C. The supernatant was collected and filtered through a 0.45 µm syringe filter (Millipore, Bayswater, Australia), then subjected to protein purification using QIAGEN Ni-NTA agarose (Qiagen, Melbourne, Australia) according to the manufacturer’s instructions. Endotoxins were removed from the purified protein using Pierce™ High Capacity Endotoxin Removal Spin Columns (Thermo Fisher Scientific). The correct size of purified Cas9 proteins was confirmed on a NuPAGE 12% Bis-Tris Gel (Thermo Fisher Scientific) and concentration was determined using the Bradford assay [42].

### 2.4. *In vitro* DNA cleavage assay

To test the enzymatic activity of expressed Cas9 variants, we conducted *in vitro* digestion of PCR-amplified *Sry* DNA with the nucleases. The *Sry* amplicon was amplified from bovine genomic DNA using primer pair *Sry*-F and *Sry*-R (**Figure 2A**, **Supplementary Table S1**), then purified using a QIAquick Gel Extraction Kit (Qiagen) according to the manufacturer’s instructions. Ribonucleoprotein (RNP) complexes were produced by incubating sgRNAs (30 nM) and Cas9 nuclease (30 nM) in NEBuffer r3.1 (New England Biolabs) at room temperature for 10 min. The purified PCR product (3 nM) was then mixed with the RNP complexes and further incubated for 15 min at 37°C. Cleavage reactions were stopped by adding 1 μl Proteinase K (20 mg/ml) (Qiagen) and followed by incubation at room temperature for 10 min. The cleaved DNA samples were analyzed by 1.8% agarose gel electrophoresis.

**Figure 2.**
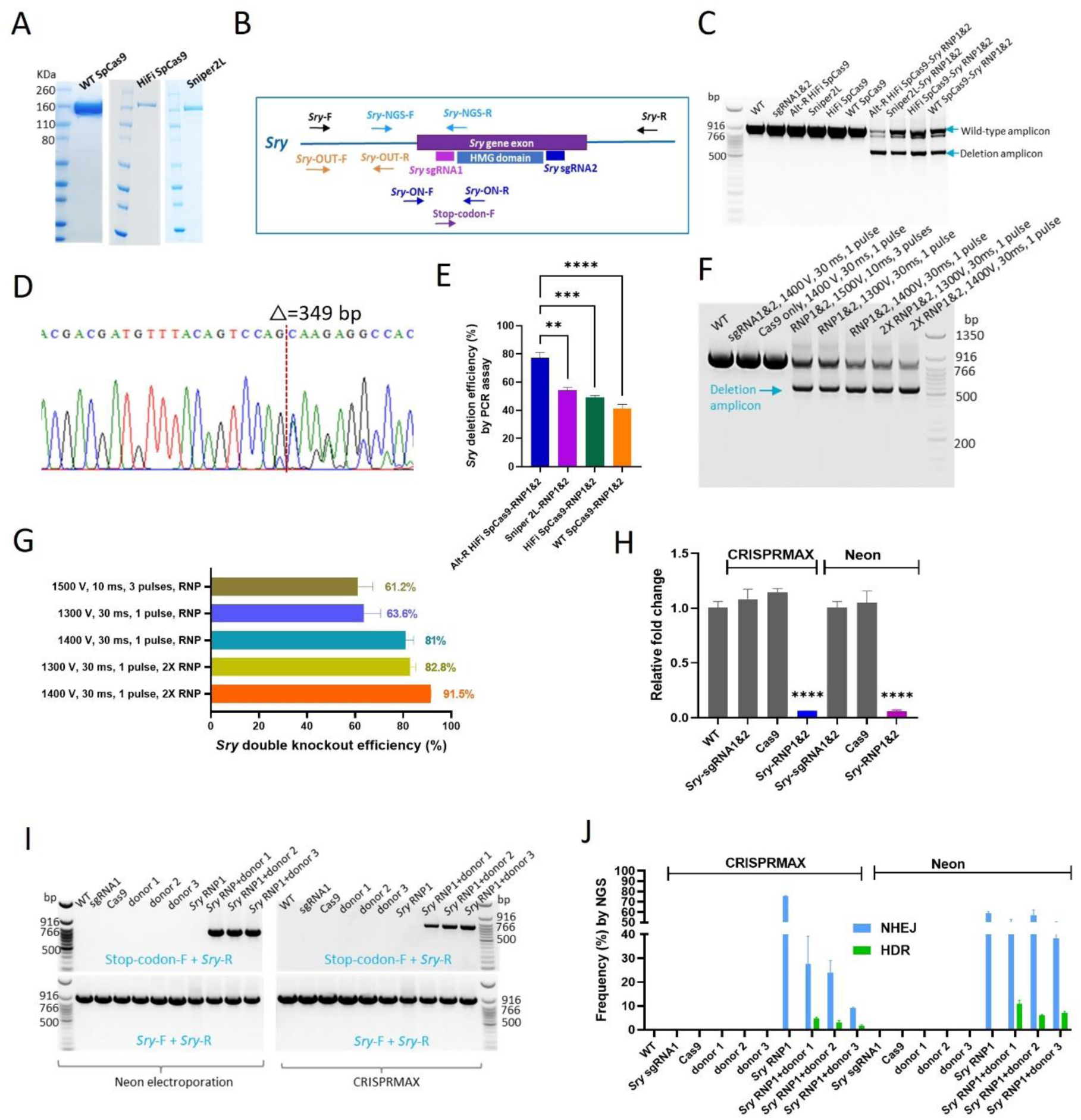
CRISPR/Cas9-mediated editing of *Sry* in MDBK cells. (**A**) Purified wild-type (WT) SpCas9, HiFi SpCas9 and Sniper2L detected in SDS-PAGE gels. (**B**) Schematic diagram showing locations of primers used for analysis of *Sry* editing. (**C**) PCR amplicons visualized in 1.8% agarose gel showing deletions of *Sry* fragment in MDBK cells with different Cas9 variants. Genomic DNA extracted from untreated WT cells and cells transfected with *Sry*-sgRNA1&2, Alt-R HiFi SpCas9, Sniper2L, HiFi SpCas9, WT SpCas9, Alt-R HiFi SpCas9-*Sry* RNP 1&2, Sniper2L-*Sry* RNP 1&2, HiFi SpCas9-*Sry* RNP 1&2 or WT SpCas9-*Sry* RNP 1&2 were used as PCR templates. (**D**) Sanger sequencing analysis showing the sequence of the deletion amplicons. (**E**) Efficiency of *Sry* deletion in MDBK cells mediated by different Cas9 variants transfected with CRISPRMAX, determined by quantification of the PCR band densities (A) using FIJI ImageJ. Mean values for three replicate experiments are shown. (**F**) PCR assays demonstrating *Sry* deletion in MDBK cells by Alt-R HiFi SpCas9 RNPs delivered with different Neon electroporation parameters. Genomic DNA obtained from untreated WT cells and cells electroporated with different parameters were subjected to PCR assays. (**G**) Efficiency of *Sry* deletion for different Neon electroporation conditions, determined by quantification of PCR band densities in (F) with FIJI ImageJ software. Mean values for duplicate experiments are shown. (**H**) Real-time PCR quantification of *Sry* deletion efficiency induced by RNPs delivered by the two different approaches. Genomic DNA was extracted and amplified from WT cells, cells CRISPRMAX (CMAX)-transfected with *Sry* sgRNA1&2, Cas9, or *Sry* RNP1&2, or cells Neon-electroporated with *Sry* sgRNA1&2, Cas9, or *Sry*RNP 1&2. The real-time PCR assays were performed in triplicate. (**I**) PCR analysis revealing site-specific integration of the transgene into the *Sry* gene in MDBK cells, using Neon electroporation or CRISPRMAX transfection for RNP and donor DNA delivery. Transgene integration at the *Sry*-sgRNA1 cleavage site was evidenced by a transgene-specific product (709 bp) amplified with Stop-codon-F and *Sry*-R primers, observed only for cells treated with both *Sry*-RNP1 and donor template 1, 2 or 3. A positive control PCR product (898 bp) amplified using the *Sry*-F and *Sry*-R primers was observed in all groups. (**J**) Frequency of NHEJ and HDR events in *Sry* knockout experiments (RNP1 only) and knock-in experiments (RNP1 + donor DNA template), determined by amplicon deep sequencing. Two biological replicates were used for *Sry*-edited groups. (** *p* value<0.01, *** *p* value<0.001, **** *p* value<0.0001, One-way ANOVA).

### 2.5. CRISPRMAX transfection of MDBK cells

MDBK cells were cultivated at 37°C in 5% O_2_ in DMEM/F-12 (Thermo Fisher Scientific) medium supplemented with 10% (v/v) fetal bovine serum (FBS, Bovogen, Melbourne, Australia), 1× GlutaMAX Supplement (Thermo Fisher Scientific) and 1× Antibiotic-Antimycotic (Thermo Fisher Scientific). All cells used in this study were below passage 12. One day prior to transfection, MDBK cells were plated onto 24-well cell culture plates (Thermo Fisher Scientific) at 1.28×10^4^ cells per well, so that the cells reached 30-70% confluence by the time of transfection. After 24 h, 1 µg Cas9 nuclease, 200 ng sgRNA1 (or 100 ng sgRNA1 and 100 ng sgRNA2) and 2 μl Cas9 Plus Reagent (Thermo Fisher Scientific) were added into 25 μl of Opti-MEM medium (Thermo Fisher Scientific) and incubated at room temperature for 5 min to form the RNP complex [43]. For knock-in experiments, 500 ng HDR donor template was also added into the Cas9 RNP solution. This mixture was then added into 25 μl of Opti-MEM medium containing 1.5 μl CRISPRMAX Transfection Reagent (Thermo Fisher Scientific) and inoculated for 15 min at room temperature before adding to the cell culture. Cells were further cultivated at 37°C in 5% CO_2_ and harvested two days post transfection.

### 2.6. Neon electroporation of MDBK cells

Electroporation was performed using the Neon Transfection System (MPK5000) (Thermo Fisher Scientific) according to the manufacturer’s instructions. Briefly, 1 µg Cas9 nuclease and 200 ng sgRNA1 (or 100 ng sgRNA1 and 100 ng sgRNA2) with/without 500 ng HDR donor template were added to 10 μl Resuspension Buffer R (Thermo Fisher Scientific) followed by vortexing and incubation at RT for 5 min to form Cas9 RNP complexes. Subsequently, 10^5^ MDBK cells were resuspended in the prepared Cas9 RNP buffer, transferred into 10 μl Neon tips, then subjected to electroporation. The electroporated cells were immediately transferred to a 24-Well Cell Culture Plate containing 500 µl DMEM/F-12 medium plus 10% (v/v) FBS and 1× GlutaMAX Supplement and cultivated at 37°C in 5% CO_2_ for 48 h before collection for analysis.

### 2.7. Screening for gene deletion and knock-in mutants by PCR assays

Genomic DNA of transfected cells was extracted using DNeasy Blood & Tissue Kit (Qiagen) as per the manufacturer’s instructions. PCR assays were performed using extracted genomic DNA as template and deletion screening primer pairs *Sry-*F + *Sry*-R, *PRLR*-F + *PRLR*-R and *Nanos2*-F + *Nanos2-R* for evaluating deletions in *Sry*, *PRLR* and *Nanos2*, respectively (**Figures 2A, 3A, 4A; Supplementary Table S1**). Genomic DNA extracted from cells in knock-in experiments was also subjected to PCR assays employing a transgene-specific forward primer (Stop-codon-F) paired with a reverse primer (*Sry*-R, *PRLR*-R or *Nanos2*-R) to detect the integration of the relevant transgenes. Each PCR reaction (25 µl) contained 12.5 µl NEBNext® High-Fidelity PCR Master Mix (New England Biolabs) with 400 nM of each primer and 50 ng template genomic DNA. The PCR program was set as follows: 98°C for 2 min, then 35 cycles of 98°C for 30 s, 65°C for 30 s and 72°C for 15 s, with a final extension at 72°C for 2 min. PCR products were analyzed using agarose gel electrophoresis and the intensity of PCR bands was quantified with FIJI/ImageJ software [44]. PCR amplicons of the expected size were excised from the gel and purified using Monarch® DNA Gel Extraction Kit (New England Biolabs). Purified PCR amplicons were subjected to Sanger sequencing carried out by The Australian Genome Research Facility (AGRF, Brisbane, Australia).

**Figure 3.**
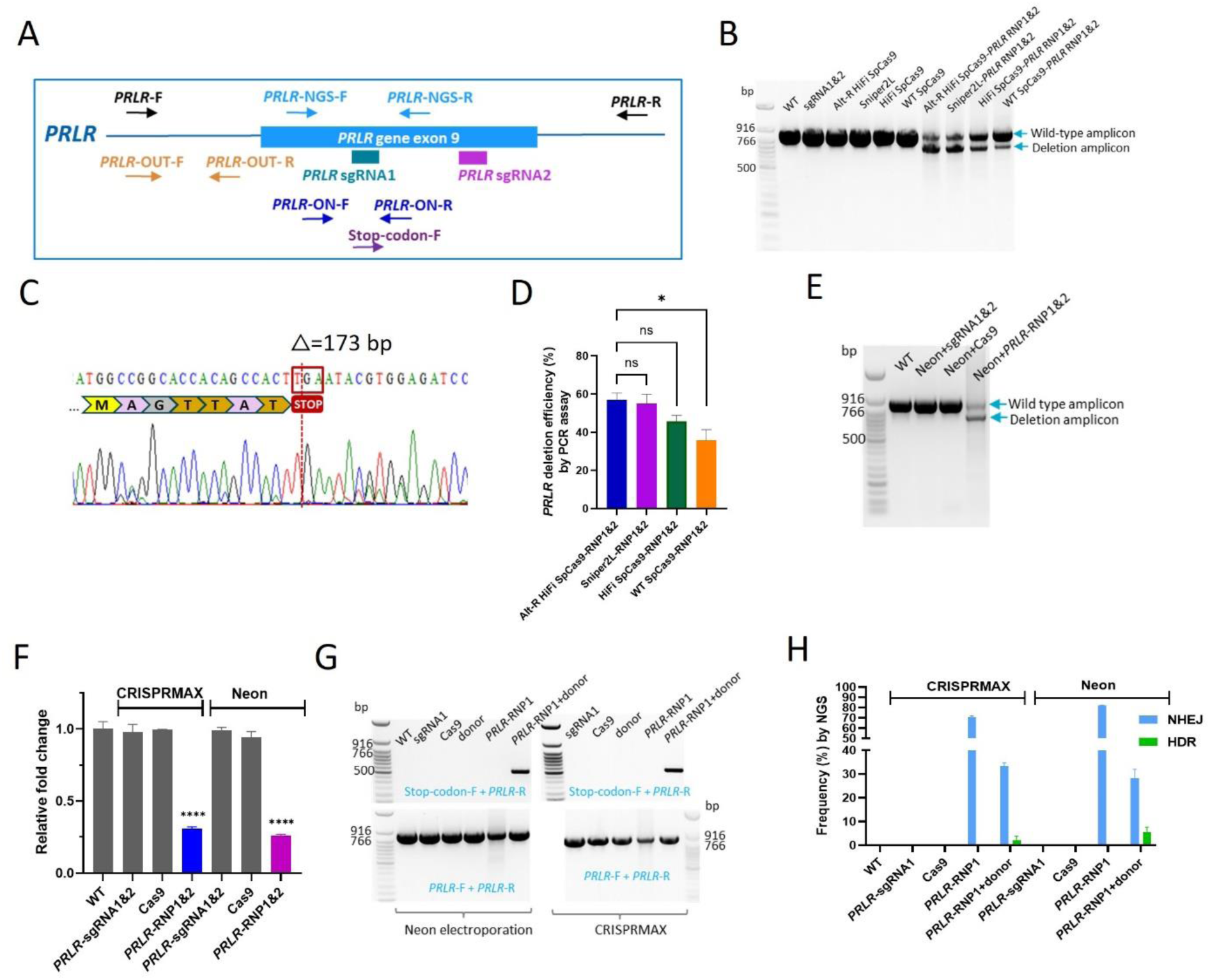
CRISPR/Cas9 editing of *PRLR* in MDBK cells. (**A**) Binding sites and directionality of primers used for assessing *PRLR* editing. (**B**) PCR assays showing deletions in the *PRLR* gene in MDBK cells, using four Cas9 variants transfected with CRISPRMAX. PCR products were amplified from genomic DNA obtained from WT untreated cells or cells treated with *PRLR*-sgRNA1&2 only, Cas9 protein only (Alt-R HiFi SpCas9, Sniper2L, HiFi SpCas9, or WT SpCas9), or *PRLR* sgRNAs1&2 complexed with one of the four Cas9 protein variants (RNPs). (**C**) Sanger sequencing chromatogram confirming a 173 bp deletion in the *PRLR* gene between the sgRNA 1&2 cleavage sites. (**D**) Efficiency of *PRLR* deletion in MDBK cells using different Cas9 variant RNPs transfected with CRISPRMAX, determined by quantification of the PCR band densities in (B) using FIJI ImageJ software. Mean values for triplicate experiments are shown. (**E**) PCR assays showing *PRLR* deletion in MDBK cells using Alt-R HiFi SpCas9 RNPs delivered by Neon electroporation. Cells were untreated or electroporated with sgRNA1&2 only, Alt-R HiFi SpCas9 protein only, or Alt-R HiFi SpCas9-*PRLR*-RNP1&2. (**F**) Quantification of *PRLR* deletion efficiency using real-time PCR assays (in triplicate), comparing the two delivery approaches. Genomic DNA was extracted and amplified from WT cells, cells CRISPRMAX-transfected with *PRLR* sgRNA1&2, Cas9, or *PRLR* RNP1&2, or cells Neon-electroporated with *PRLR* sgRNA1&2, Cas9, or *PRLR* RNP1&2. The real-time PCR assays were carried out in triplicates. (**G**) PCR products demonstrating site-specific knock-in of the *PRLR* transgene, using Neon electroporation or CRISPRMAX transfection to delivery RNP and donor DNA template. Successful knock-in to the *PRLR* gene was indicated by a transgene specific amplicon (510 bp) observed only for cells treated with both *PRLR*-RNP1 and *PRLR*-donor, delivered with CRISPRMAX or Neon electroporation. A positive control amplicon (859 bp) was detected in all treatments. (**H**) NGS determining the frequency of NHEJ and HDR events in *PRLR* knockout and knock-in groups. Two biological duplicates were performed for *PRLR*-edited groups. (ns, not significant, * *p* value<0.05, ** *p* value<0.01, One-way ANOVA).

**Figure 4.**
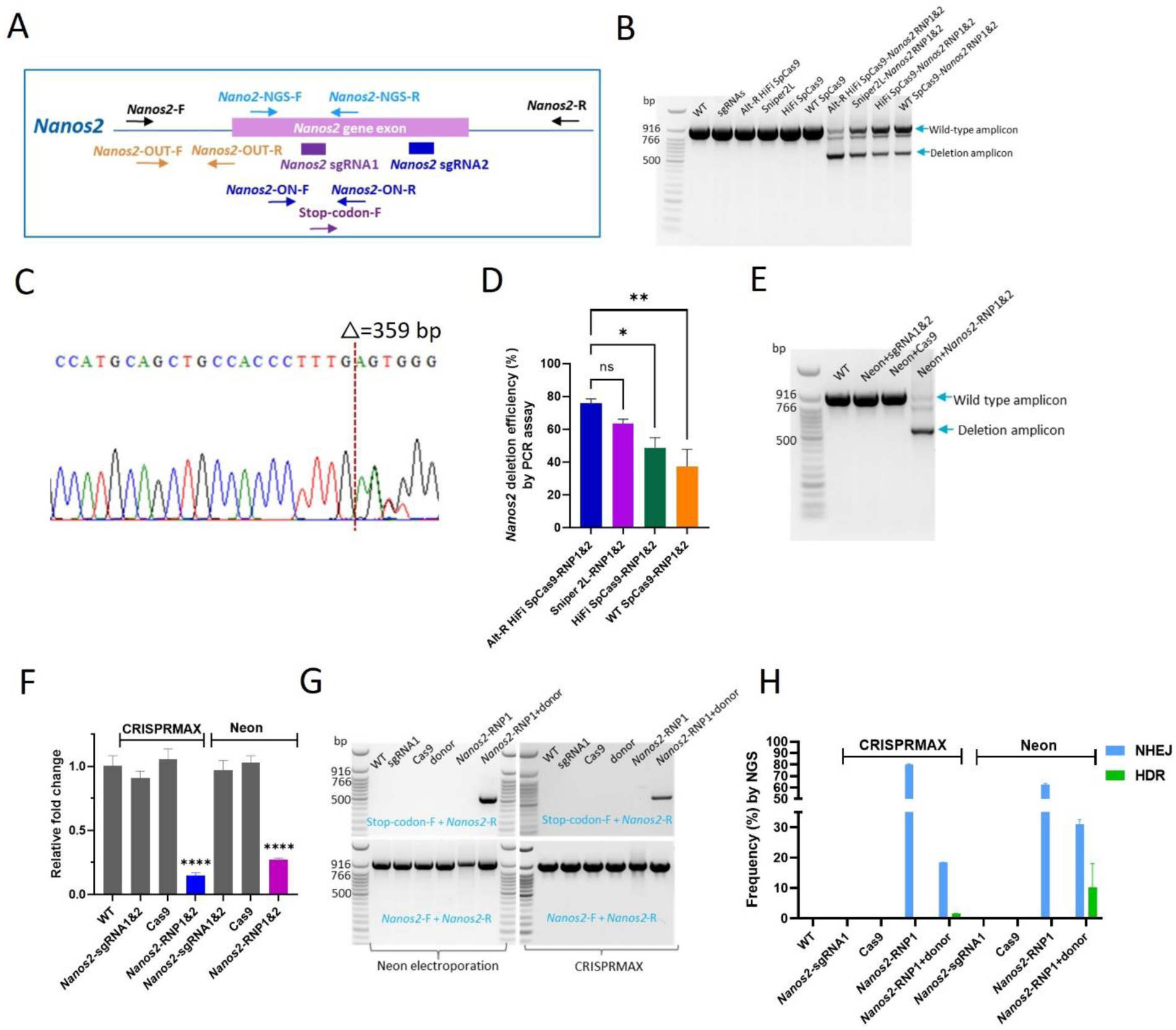
CRISPR/Cas9 editing of *Nanos2* in MDBK cells. (**A**) Binding sites and directionality of primers used for investigating *Nanos2* editing. (**B**) PCR products demonstrating deletion in the *Nanos2* gene in MDBK cells, using the four Cas9 protein variants transfected with CRISPRMAX. PCR products were amplified from genomic DNA extracted from WT cells or cells treated with *Nanos2*-sgRNA1&2 only, Cas9 protein only (Alt-R HiFi SpCas9, Sniper2L, HiFi SpCas9, or WT SpCas9), or *Nanos2* sgRNAs1&2 complexed with one of the four Cas9 protein variants (RNPs). (**C**) Sanger sequencing chromatogram confirming a 359 bp deletion in the *Nanos2* gene between the sgRNA1&2 cleavage sites. (**D**) Efficiency of *Nanos2* deletion in MDBK cells using different Cas9 variant RNPs transfected with CRISPRMAX, determined by quantification of the PCR product densities in (B) with FIJI ImageJ. The experiment was performed in triplicates. (**E**) PCR amplicons showing *Nanos2* deletion in MDBK cells by Alt-R HiFi SpCas9 RNPs delivered by Neon electroporation. Cells were untreated or electroporated with *Nanos2*-sgRNA1&2 only, Alt-R HiFi SpCas9 protein only, or Alt-R HiFi SpCas9-*Nanos2*-RNP1&2. (**F**) Quantification of *Nanos2* deletion efficiency using real-time PCR assays (performed in triplicate), comparing the two RNP delivery methods. Genomic DNA was extracted and amplified from WT cells, cells CRISPRMAX-transfected with *Nanos2-*sgRNA1&2, Cas9, or *Nanos2* RNP1&2, or cells Neon-electroporated with *Nanos2*-sgRNA1&2, Cas9, or *Nanos2* RNP1&2. (**G**) PCR detection of site-specific integration of the *Nanos2* transgene. A transgene-specific amplicon (509 bp) was observed only in cells treated with both *Nanos2*-RNP1 and *Nanos2*-donor, delivered by CRISPRMAX or Neon electroporation. A positive control amplicon (910 bp) was seen in all groups. (**H**) Amplicon NGS illustrating the frequency of NHEJ and HDR events in *Nanos2* knockout and knock-in experiments. Two biological repeats were used for *Nanos2*-edited groups. (ns, not significant, * *p* value<0.05, ** *p* value<0.01, **** *p* value<0.0001, One-way ANOVA).

### 2.8. Evaluating gene deletion efficiency using real-time PCR

Quantitative real-time PCR (qPCR) was conducted to assess deletion efficiency at the target genes [45,46] using template genomic DNA extracted from CRISPRMAX or Neon electroporation transfected cells. Each genomic DNA template was subjected to two qPCR reactions using ‘ON’ primers and ‘OUT’ primers (**Figures 2A, 3A, 4A, Supplementary Table S1**). The ‘OUT’ (flanking) primer pair (OUT-F + OUT-R) was designed to amplify a control PCR product (for normalization purposes) from sequence at least 50 bp distant to the sgRNA targeting site. The ‘ON’ (overlapping) primer pair included a forward primer ON-F and a reverse ON-R primer which located in the region between the targeting loci of sgRNA1 and sgRNA2; these primers could amplify a product from wild-type allelic copies but not from allelic copies with the expected sequence deletion. This method uses the fact that gene editing causes a deletion at the target locus that prevents binding and amplification with the ON primers; the reduction in wild-type allele copies in the template DNA results in delayed real-time PCR amplification with the ON primers and a higher quantification cycle (Cq) value [45,46]. Binding and amplification with the OUT primers located outside the deleted sequence are unaffected. Accordingly, the gene editing efficiency can be calculated by comparison of the ON/OUT Cq ratios of edited and non-edited groups [45,46]. The qPCR assays were performed using the ViiA 7 Real-Time PCR System (Applied Biosystems, Waltham, Massachusetts, USA) with 384-well block. Each 7 ul qPCR reaction (performed in triplicate) included 3.5 µl SYBR Green Universal Master Mix (Applied Biosystems), 25 ng genomic DNA, and 0.4 µM of each primer. The thermal cycling parameters were 95°C for 10 min, 40 cycles of 95°C for 30 s and 60°C for 30 s, followed by 72°C for 30 s.

### 2.9. Amplicon deep sequencing

Amplicon-based NGS was conducted to assess the NHEJ and HDR outcomes in gene-edited cells. For each of the three genes, amplicon primer pairs with Nextera 5’ tags (**Supplementary Table S1**) were designed to flank the sgRNA1 cleavage site (**Figures 2A, 3A, 4A; Supplementary Table S1**). Genomic DNA was extracted from treated cells and amplified with the relevant amplicon primer pair. Each PCR reaction (25 µl) included 12.5 µl NEBNext Ultra II Q5 Master Mix (New England Biolabs), 30 ng genomic DNA, 1 µM of each primer and Nuclease-Free H_2_O (ThermoFisher Scientific). Thermal cycling involved denaturation at 98°C for 30 s, 35 cycles of 98°C for 10 s, 68°C for 30 s and 72°C for 30 s, and a final extension for 2 min at 72°C. PCR amplicons (200 bp, 211 bp and 213 bp for *Sry*, *PRLR* and *Nanos2*, respectively) obtained from three independent PCR reactions for each genomic DNA extract were pooled to generate uniquely indexed libraries. The libraries were then subjected to 150 bp paired-end sequencing on the Illumina MiSeq platform at AGRF. Fastq files were analyzed with CRISPResso2 to determine on-target gene editing efficiency [47].

### 2.10. Statistical analysis

All statistical analyses were performed with GraphPad Prism software (Version 9.1.2, La Jolla, CA, USA). Statistical significance was analyzed using One-way ANOVA followed by Dunnett’s multiple comparisons test. Differences for particular comparisons were considered to be statistically significant where *p* < 0.05. * *p* value < 0.05, ** *p* value < 0.01, *** *p* value < 0.001, **** *p* value < 0.0001.

## 3.1. Results

### 3.1. Expression and purification of Cas9 variants

Three recombinant SpCas9 protein variants: Sniper2L [38], HiFi SpCas9 [37], and SpCas9 [36] were expressed in *E. coli* BL21(DE3) and purified with Ni-NTA affinity chromatography. SDS-PAGE analysis verified that purified Sniper2L, HiFi SpCas9 and WT SpCas9 all had the predicted size of approximately 164 kDa (**Figure 2A**). The enzymatic activity of generated Cas9 nucleases was confirmed by *in vitro* DNA cleavage assay and notably, all three in-house produced Cas9 variants demonstrated *in vitro* cleavage efficiency similar to the commercial Alt-R HiFi SpCas9 (**Supplementary Figure S1**).

### 3.2. Comparing the efficiency of different Cas9 variants in MDBK cells

To compare the gene editing efficacy of the three produced Cas9 nucleases with the IDT Alt-R HiFi SpCas9 in MDBK cells, we performed targeted gene deletion by lipofectamine CRISPRMAX transfection of RNP complexes [43], using two sgRNAs for each target gene. PCR assays were conducted using deletion screening primer pairs (covering the dual target cleavage sites for each gene. *Sry* WT PCR bands (898 bp) amplified with primers *Sry*-F and *Sry*-R (**Figure 2B**, **Supplementary Table S1**) were observed in both control groups (untreated cells, cells treated with *Sry*-sgRNA1&2 only, or cells treated with one of the four nucleases without sgRNAs) and edited groups (cells treated with one of the four nucleases as RNP complexed with *Sry-*sgRNA1&2) (**Figure 2C**). Deletion amplicons with the expected size of 548 bp were only detected in the *Sry*-edited groups. The deletion amplicons were subjected to Sanger sequencing, which confirmed the deletion of 349 bp gene fragment including the HMG domain in *Sry* (**Figure 2D**). Deletion efficiency was measured by quantifying the density of PCR bands in Figure 2C using FIJI ImageJ software. For *Sry*, Alt-R HiFi SpCas9 demonstrated the highest deletion efficiency (77%), followed by Sniper2L (54%), HiFi SpCas9 (49%) and WT SpCas9 (41%) (**Figure 2E**).

Similarly, for *PRLR*, a WT PCR band generated using deletion screening primer pair *PRLR*-F and *PRLR*-R (**Figure 3A**, **Supplementary Table S1**) with expected size of 859 bp was detected in all groups whereas the deletion bands (686 bp) were only observed in *PRLR*-edited groups (cells treated with one of the four Cas9 variants complexed as RNP with *PRLR-*sgRNA1&2) (**Figure 3B**). Sanger sequencing of the *PRLR* deletion amplicons confirmed the expected deletion (173 bp) of *PRLR* (**Figure 3C**). Notably, Sniper2L (55%) exhibited comparable deletion efficiency to Alt-R HiFi SpCas9 (57%) for *PRLR.* HiFi SpCas9 (46%) showed lower (but not significantly different) efficiency whereas WT SpCas9 (36%) was significantly less efficient than the IDT nuclease (**Figure 3D**).

Likewise, for *Nanos2*, a 910 bp WT PCR band amplified using primers *Nanos2*-F and *Nanos2*-R (**Figure 4A**, **Supplementary Table S1**) was observed in all samples whilst the deletion band (551 bp) was only detectable in cells treated with one of the four Cas9 variants complexed as RNP with *Nanos2-*sgRNA1&2 (**Figure 4B**). The gene deletion (359 bp) was confirmed by Sanger sequencing (**Figure 4C**). For *Nanos2* deletion, Alt-R HiFi SpCas9 showed slightly but not significantly higher efficiency (76%) than Sniper2L (63%), whereas HiFi SpCas9 (49%) and WT SpCas9 (37%) were significantly less efficient than the IDT nuclease (**Figure 4D**).

### 3.3. Selection of Neon electroporation parameters

To optimize the Neon electroporation system in MDBK cells, using *Sry* as the target and Alt-R HiFi SpCas9, we tested the following sets of electrical parameters: 1) 1500 V, 10 ms, 3 pulses; 2) 1300 V, 30 ms, 1 pulse; 3) 1400 V, 30 ms, 1 pulse; 4) 1300 V, 30 ms, 1 pulse, 2x amount of RNP; and 5) 1400 V, 30 ms, 1 pulse, 2x amount of RNP. PCR assays were performed to assess the efficiency of *Sry* deletion for the different parameters. *Sry* deletion amplicons with expected size of 548 bp were observed in cells treated with *Sry*-RNP1&2 delivered via Neon electroporation for all the tested sets of parameters (**Figure 2F**). Densitometric quantification of the PCR bands indicated that Parameter set 5 was the most efficient for *Sry* deletion (91.5%) followed by Parameter set 4 (82.8%), Parameter set 3 (81%), Parameter set 2 (63.6%) and Parameter set 1 (61.2%) (**Figure 2G**). Neon electroporation of Alt-R HiFi SpCas9 RNPs with *PRLR*-sgRNAs1&2 or *Nanos2*-sgRNAs1&2 using Parameter set 5 also induced efficient deletion of *PRLR* (**Figure 3E**) and *Nanos2* (**Figure 4E**) in MDBK cells, respectively. Therefore, we selected these parameters for subsequent experiments involving Neon electroporation.

### 3.4. Efficient gene deletion determined by real-time PCR

To evaluate the efficiency of CRISPRMAX lipofection and Neon electroporation in delivering RNP complexes to MDBK cells, we performed qPCR assays [45,46] using genomic DNA extracted from CRISPRMAX or Neon electroporation transfected cells and two pairs of primer pairs (ON primers and OUT primers). We found that CRISPRMAX + *Sry-*RNP1&2 and Neon electroporation + *Sry-*RNP1&2 treatments of MDBK cells significantly reduced wild-type (uncut) *Sry* template DNA by 93.9% (*p* < 0.0001) and 94.1% (*p* < 0.0001), respectively (**Figure 2H**). Similarly, significant reductions in the targeted WT sequences were observed in equivalent experiments targeting *PRLR* [CRISPRMAX, 70% (*p* < 0.0001), Neon electroporation, 73.3% (*p* < 0.0001)] (**Figure 3F**), and *Nanos2* [CRISPRMAX, 85% (*p* < 0.0001), Neon electroporation, 73% (*p* < 0.0001)] (**Figure 4F**).

### 3.5. PCR assays revealed site-specific knock-ins

To investigate the efficiency of transgene integration (knock-in) at the target loci using CRISPRMAX or Neon transfection, we designed a 104 nt single-stranded DNA donor template for each target gene, with homology arms (40 nt) matching sequence flanking the sgRNA1 cleavage site for each gene. These donor oligonucleotides were transfected into MDBK cells in combination with Alt-R HiFi SpCas9 RNP incorporating the corresponding sgRNA1 for each gene. Genomic DNA from treated cells was subjected to a transgene-specific PCR to identify successful knock-in and a positive control PCR. For detection of successful knock-ins, primer Stop-codon-F was paired with primer *Sry*-R, *PRLR-*R, or *Nanos2-*R to amplify a transgene-specific amplicon from the genomic DNA of cells treated with Cas9-RNPs targeting *Sry, PRLR,* or *Nanos2*, respectively (**Figures 2A, 3A, 4A; Supplementary Table S1**). Transgene-specific amplicons with expected sizes of 709 bp (*Sry,* **Figure 2I**), 510 bp (*PRLR*, **Figure 3G**) or 509 bp (*Nanos2*, **Figure 4G**) were only observed for cells treated with the corresponding Cas9-RNP-sgRNA1 and donor oligo for each gene, for both CRISPRMAX and Neon electroporation transfection. No transgene-specific PCR amplicons were detected in the control groups; untreated, sgRNA1 only, Cas9 only, donor oligo only, or Cas9-sgRNA RNP without donor oligo. Positive control amplicons with expected sizes of 898 bp (*Sry*, **Figure 2I**), 859 bp (*PRLR*, **Figure 3G**), or 910 bp (*Nanos2*, **Figure 4G**) appeared in all corresponding treatments including controls. These results indicate the successful knock-in of transgenes at the target loci.

### 3.6. Detection of NHEJ and HDR frequency using amplicon deep sequencing

Amplicon NGS was carried out to quantify the efficiency of NHEJ and HDR in edited cells. Genomic DNA extracted from cells treated with *Sry*-RNP1 ± *Sry*-donor, *PRLR*-RNP1 ± *PRLR*-donor or *Nanos2*-RNP1 ± *Nanos2-*donor was sequenced on the Illumina MiSeq platform (150 bp paired-end), and the data were analyzed with CRISPResso2 software [47]. NGS analysis demonstrated the frequency of indels in the *Sry* knockout group (treated with *Sry*-RNP1, no donor) was higher for CRISPRMAX transfection (76%) than Neon electroporation (60%) (**Figure 2J**). For knock-in into the *Sry* gene, we evaluated three differently modified variants of the HDR donor ssODN: an Alt-R HDR modified donor (#1), a PS bond modified donor (#2) and a non-modified donor (#3). Deep sequencing revealed that for CRISPRMAX transfection, in combination with *Sry*-RNP1, donor 1 yielded 28% NHEJ and 4.8% HDR, donor 2 yielded 24% NHEJ and 3.2% HDR, and donor 3 induced 9.2% NHEJ and 1.8% HDR (**Figure 2J**). However, Neon electroporation of *Sry*-RNP1 in combination with the same donors produced superior results; donor 1 produced 46% NHEJ and 11% HDR, donor 2 generated 57% NHEJ and 6.1% HDR, and donor 3 induced 38.4% NHEJ and 7.1% HDR (**Figure 2J**).

Given that donor 1 induced the highest HDR frequency for *Sry*, we used the same Alt-R HDR modified ssODN donor type for knock-in into *PRLR* and *Nanos2*. CRISPRMAX transfection and Neon electroporation of *PRLR*-RNP1 without donor oligo produced 70.5% and 82.4% NHEJ events in treated cells, respectively (**Figure 3H**). Co-delivery of *PRLR*-RNP1 with *PRLR-*donor resulted in 33.5% NHEJ and 2.2% HDR for CRISPRMAX transfection, and 28.3% NHEJ and 5.5% HDR for Neon electroporation (**Figure 3H**). Similarly, NHEJ events in cells transfected with *Nanos2*-RNP1 without donor oligo were 80% and 63% for CRISPRMAX transfection and Neon electroporation, respectively (**Figure 4H**). CRISPRMAX transfection and Neon electroporation of cells with *Nanos2*-RNP1 and *Nanos2-*donor yielded 18.4% NHEJ and 1.5% HDR, and 31% NHEJ and 10.2% HDR, respectively (**Figure 4H**).

## 4. Discussion

CRISPR/Cas9 is a powerful genome engineering tool that can achieve targeted editing of endogenous genes and integration of exogenous genes [3,4,13,48,49]. To date, genome editing applications in livestock to enhance traits or create new biomedical research models have proven to be both feasible and valuable [1,3]. Improvements in the precision and effectiveness of gene editing will help to drive the adoption of gene editing for the rapid and focused advancement of livestock production. To further the aim of developing an efficient gene editing workflow in cattle, we evaluated the efficacy of different Cas9 variants and delivery methods in an immortalized bovine kidney cell line.

We evaluated the editing efficiency of in-house produced Sniper2L, WT SpCas9 and HiFi SpCas9 and a commercially available Alt-R HiFi SpCas9. Sniper2L is a newly developed HiFi SpCas9 variant which has been reported to induce highly specific and efficient gene editing in human HEK293T cells [38]. However, the performance of Sniper2L in bovine cells has not been previously assessed. *In vitro* DNA cleavage assay revealed the three in-house generated SpCas9 nucleases possessed similar enzymatic activity to the commercial Cas9. However, they exhibited variable efficacy in MDBK cells, as evidenced by PCR assays. Specifically, when targeting the *Sry* gene, the commercial Alt-R HiFi SpCas9 outperformed the three in-house produced Cas9 nucleases. However, Sniper2L displayed comparable efficiency to Alt-R HiFi SpCas9 when editing *PRLR* and *Nanos2*. Our finding that the Cas9 variants exhibited similar cleavage efficiency *in vitro* but demonstrated variable editing efficiency in cells may be partially explained by a difference in the nuclear localization signal (NLS), which has been shown to impact Cas9 performance [50–52]. The three in-house generated Cas9 variants have 3× NLS signals fused to the C-terminal end of the protein, whereas the commercial Alt-R HiFi SpCas9 has a proprietary, unspecified NLS configuration. It seems likely that this configuration contributed to the superior efficiency of Alt-R HiFi SpCas9 in the cellular environment. Further modifications to the NLS configuration of the in-house produced Cas9 proteins might therefore enhance their performance. In addition, we identified that Sniper2L may be an excellent choice for precise gene editing in cattle, as it exhibited editing efficiency close to the Alt-R HiFi Cas9 in some instances.

Next, we compared the efficiency of two CRISPR/Cas9 delivery approaches, Lipofectamine CRISPRMAX transfection and Neon electroporation [53–56], for delivering RNPs and donor templates into MDBK cells. Selection between these two methods will generally depend on the specific requirements of the experiment, such as the type of cells and the desired efficiency of gene editing. Lipofectamine CRISPRMAX offers ease of use with low cell toxicity and it is a more cost-effective option than the Neon system [43]. The Neon is superior at transfecting hard-to-transfect cells [43] but does incur higher costs (although a published protocol is available to regenerate the expensive Neon tips [57]). In this study, we compared the efficiency of the two delivery methods for performing targeted gene knockout and knock-in in MDBK cells. Real-time PCR assays [45,46] showed that the deletion frequencies generated through CRISPRMAX transfection and Neon electroporation were comparable for all three target genes, ranging from 70% to 94%. Deep sequencing of amplicons produced using Nextera-tagged and target-specific primers also confirmed that both delivery approaches achieved a high frequency (60-82%) of DNA cleavage repaired by NHEJ for all three target genes. However, the frequency of precise knock-in of a ssODN donor template by HDR was ≤11% for both delivery methods, with Neon electroporation (5.5-11%) proving more effective than CRISPRMAX (1.5-4.8%). Additionally, the NGS analysis revealed a high frequency of NHEJ events (9.8-57%) in the knock-in groups.

It is known that there is competition between the NHEJ and HDR pathways in the repair of DSBs in DNA [58], and mammalian cells preferentially employ NHEJ over HDR [59,60]. The frequency of HDR in MDBK cells might be further improved by strategies that inhibit critical factors in the NHEJ pathway (e.g. a small-molecule inhibitor, SCR7) [61–64] while enhancing the HDR event (e.g. a HDR enhancer, RS-1) [65–68]. Previous studies have also shown that HDR efficiency can be increased by capturing cells in the S and G2 phases at the time of delivery of CRISPR components, when the HDR pathway is most active [69]. However, these approaches may negatively affect cell viability, which need to be carefully considered [69]. In addition, HDR frequency might also be improved through optimizing the components in the CRISPR machinery. For example, Kato-Inui *et al*. utilized modified sgRNAs and Cas9 variants with higher conformational checkpoint threshold and achieved significantly enhanced the HDR efficiency on both single-nucleotide editing and long fragment (multi-kb) integration in human cells [70]. Furthermore, alternative methods such as Homology-Independent Targeted Integration (HITI), which takes advantage of the highly efficient NHEJ pathway and enables a high frequency of knock-ins in both dividing and non-dividing cells [71], could presumably be used to enhance the efficiency of precise transgene integrations into MDBK cells.

## 5. Conclusion

In conclusion, our results showed the commercial Alt-R HiFi SpCas9 excelled in *Sry* editing, while Sniper2L, an engineered high-fidelity SpCas9 variant, demonstrated comparable efficiency when editing two other genes (*PRLR* and *Nanos2*). Sniper2L may be an excellent choice for gene-editing of bovine cells when both high efficiency and high specificity are important considerations. Lipofectamine CRISPRMAX and Neon electroporation delivery induced comparable NHEJ frequencies in MDBK cells, but Neon electroporation was superior for HDR-mediated knock-in of ssODN donor template. These findings provide valuable insights for optimizing genome editing outcomes in cattle and may help to promote accelerated and widespread application of gene engineering technology in livestock.

## CRediT authorship contribution statement

**Xiaofeng Du:** Writing – review & editing, Writing – original draft, Visualization, Validation, Methodology, Investigation, Data curation, Conceptualization. **Alexander Quinn:** Writing – review & editing, Writing – original draft, Validation, Methodology. **Timothy Mahony:** Writing – review & editing, Visualization, Resources. **Di Xia:** Writing – review & editing, Methodology, Resources. **Laercio Porto-Neto:** Writing – review & editing, Investigation, Supervision, Resources, Project administration, Funding acquisition, Conceptualization.

## Acknowledgements

We would like to thank the Australian Genome Research Facility for the technical support in preparing samples for next generation sequencing and Dr James Kijas for providing valuable comments and suggestions in revision of this paper.

## Data availability

Data will be made available on request.

## References

1. A. Menchaca, P. dos Santos-Neto, A. Mulet, M. Crispo. CRISPR in livestock: From editing to printing. Theriogenology, 150 (2020), pp. 247–254.

2. B. Petersen. Basics of genome editing technology and its application in livestock species. Reproduction in Domestic Animals, 52 (2017), pp. 4–13.

3. A. Jabbar, F. Zulfiqar, M. Mahnoor, N. Mushtaq, M. H. Zaman, A. S. U. Din, M. A. Khan, H. I. Ahmad. Advances and Perspectives in the Application of CRISPR-Cas9 in Livestock. Molecular Biotechnology, 63 (2021), pp. 757–767.

4. H. Ledford, E. Callaway. Pioneers of CRISPR gene editing win chemistry Nobel. Nature, 586 (2020), pp. 346–347.

5. R. Barrangou, J. A. Doudna. Applications of CRISPR technologies in research and beyond. Nature biotechnology, 34 (2016), pp. 933–941.

6. H. Wang, M. La Russa, L. S. Qi. CRISPR/Cas9 in genome editing and beyond. Annual review of biochemistry, 85 (2016), pp. 227–264.

7. R. Peng, G. Lin, J. Li. Potential pitfalls of CRISPR/Cas9-mediated genome editing. The FEBS journal, 283 (2016), pp. 1218–1231.

8. J. A. Doudna, E. Charpentier. The new frontier of genome engineering with CRISPR-Cas9. Science, 346 (2014), pp. 1258096.

9. J. Y. Wang, J. A. Doudna. CRISPR technology: A decade of genome editing is only the beginning. Science, 379 (2023), pp. eadd8643.

10. D. J. Dickinson, B. Goldstein. CRISPR-based methods for Caenorhabditis elegans genome engineering. Genetics, 202 (2016), pp. 885–901.

11. M. D. Szczelkun, M. S. Tikhomirova, T. Sinkunas, G. Gasiunas, T. Karvelis, P. Pschera, V. Siksnys, R. Seidel. Direct observation of R-loop formation by single RNA-guided Cas9 and Cascade effector complexes. Proceedings of the National Academy of Sciences, 111 (2014), pp. 9798–9803.

12. M. Adli. The CRISPR tool kit for genome editing and beyond. Nature communications, 9 (2018), pp. 1911.

13. L. Cong, F. A. Ran, D. Cox, S. Lin, R. Barretto, N. Habib, P. D. Hsu, X. Wu, W. Jiang, L. A. Marraffini. Multiplex genome engineering using CRISPR/Cas systems. Science, 339 (2013), pp. 819–823.

14. F. Ran, P. D. Hsu, J. Wright, V. Agarwala, D. A. Scott, F. Zhang. Genome engineering using the CRISPR-Cas9 system. Nature protocols, 8 (2013), pp. 2281–2308.

15. T. Hai, F. Teng, R. Guo, W. Li, Q. Zhou. One-step generation of knockout pigs by zygote injection of CRISPR/Cas system. Cell research, 24 (2014), pp. 372–375.

16. K.-E. Park, A. V. Kaucher, A. Powell, M. S. Waqas, S. E. Sandmaier, M. J. Oatley, C.-H. Park, A. Tibary, D. M. Donovan, L. A. Blomberg. Generation of germline ablated male pigs by CRISPR/Cas9 editing of the NANOS2 gene. Scientific Reports, 7 (2017), pp. 40176.

17. D. Miao, M. I. Giassetti, M. Ciccarelli, B. Lopez-Biladeau, J. M. Oatley. Simplified pipelines for genetic engineering of mammalian embryos by CRISPR-Cas9 electroporation. Biology of reproduction, 101 (2019), pp. 177–187.

18. S. Navarro-Serna, M. Vilarino, I. Park, J. Gadea, P. J. Ross. Livestock gene editing by one-step embryo manipulation. Journal of equine veterinary science, 89 (2020), pp. 103025.

19. W. Ni, J. Qiao, S. Hu, X. Zhao, M. Regouski, M. Yang, I. A. Polejaeva, C. Chen. Efficient gene knockout in goats using CRISPR/Cas9 system. PloS one, 9 (2014), pp. e106718.

20. Y. Yin, H. Hao, X. Xu, L. Shen, W. Wu, J. Zhang, Q. Li. Generation of an MC3R knock-out pig by CRSPR/Cas9 combined with somatic cell nuclear transfer (SCNT) technology. Lipids in health and disease, 18 (2019), pp. 1–8.

21. F. Schuster, P. Aldag, A. Frenzel, K.-G. Hadeler, A. Lucas-Hahn, H. Niemann, B. Petersen. CRISPR/Cas12a mediated knock-in of the Polled Celtic variant to produce a polled genotype in dairy cattle. Scientific reports, 10 (2020), pp. 13570.

22. C. Galli, G. Lazzari. 25th ANNIVERSARY OF CLONING BY SOMATIC-CELL NUCLEAR TRANSFER: Current applications of SCNT in advanced breeding and genome editing in livestock. Reproduction, 162 (2021), pp. F23–F32.

23. G. Laible, J. Wei, S. Wagner. Improving livestock for agriculture–technological progress from random transgenesis to precision genome editing heralds a new era. Biotechnology journal, 10 (2015), pp. 109–120.

24. J. Popova, V. Bets, E. Kozhevnikova. Perspectives in Genome-Editing Techniques for Livestock. Animals, 13 (2023), pp. 2580.

25. K. Lee, K. Uh, K. Farrell. Current progress of genome editing in livestock. Theriogenology, 150 (2020), pp. 229–235.

26. J. C. Lin, A. L. Van Eenennaam. Electroporation-mediated genome editing of livestock zygotes. Frontiers in genetics, 12 (2021), pp. 648482.

27. L. S. A. Camargo, J. R. Owen, A. L. Van Eenennaam, P. J. Ross. Efficient one-step knockout by electroporation of ribonucleoproteins into zona-intact bovine embryos. Frontiers in genetics, 11 (2020), pp. 570069.

28. M. Vilarino, F. P. Suchy, S. T. Rashid, H. Lindsay, J. Reyes, B. R. McNabb, T. van der Meulen, M. O. Huising, H. Nakauchi, P. J. Ross. Mosaicism diminishes the value of pre-implantation embryo biopsies for detecting CRISPR/Cas9 induced mutations in sheep. Transgenic research, 27 (2018), pp. 525–537.

29. K. M. Whitworth, R. S. Prather. Somatic cell nuclear transfer efficiency: how can it be improved through nuclear remodeling and reprogramming? Molecular reproduction and development, 77 (2010), pp. 1001–1015.

30. K. Srirattana, M. Kaneda, R. Parnpai. Strategies to improve the efficiency of somatic cell nuclear transfer. International Journal of Molecular Sciences, 23 (2022), pp. 1969.

31. S. Madin, N. Darby Jr. Established kidney cell lines of normal adult bovine and ovine origin. Proceedings of the Society for Experimental Biology and Medicine, 98 (1958), pp. 574–576.

32. B. Li, C. Zeng, Y. Dong. Design and assessment of engineered CRISPR–Cpf1 and its use for genome editing. Nature protocols, 13 (2018), pp. 899–914.

33. L. R. Porto-Neto, D. M. Bickhart, A. J. Landaeta-Hernandez, Y. T. Utsunomiya, M. Pagan, E. Jimenez, P. J. Hansen, S. Dikmen, S. G. Schroeder, E.-S. Kim. Convergent evolution of slick coat in cattle through truncation mutations in the prolactin receptor. Frontiers in Genetics, 9 (2018), pp. 57.

34. M. D. Littlejohn, K. M. Henty, K. Tiplady, T. Johnson, C. Harland, T. Lopdell, R. G. Sherlock, W. Li, S. D. Lukefahr, B. C. Shanks. Functionally reciprocal mutations of the prolactin signalling pathway define hairy and slick cattle. Nature communications, 5 (2014), pp. 5861.

35. J. A. Zuris, D. B. Thompson, Y. Shu, J. P. Guilinger, J. L. Bessen, J. H. Hu, M. L. Maeder, J. K. Joung, Z.-Y. Chen, D. R. Liu. Cationic lipid-mediated delivery of proteins enables efficient protein-based genome editing in vitro and in vivo. Nature biotechnology, 33 (2015), pp. 73–80.

36. N. E. Sanjana, O. Shalem, F. Zhang. Improved vectors and genome-wide libraries for CRISPR screening. Nature methods, 11 (2014), pp. 783–784.

37. M. Escobar, J. Li, A. Patel, S. Liu, Q. Xu, I. B. Hilton. Quantification of genome editing and transcriptional control capabilities reveals hierarchies among diverse CRISPR/Cas systems in human cells. ACS Synthetic Biology, 11 (2022), pp. 3239–3250.

38. Y.-h. Kim, N. Kim, I. Okafor, S. Choi, S. Min, J. Lee, S.-M. Bae, K. Choi, J. Choi, V. Harihar. Sniper2L is a high-fidelity Cas9 variant with high activity. Nature chemical biology, 19 (2023), pp. 972–980.

39. J.-W. Wang, A. Wang, K. Li, B. Wang, S. Jin, M. Reiser, R. F. Lockey. CRISPR/Cas9 nuclease cleavage combined with Gibson assembly for seamless cloning. Biotechniques, 58 (2015), pp. 161–170.

40. K. Luk, P. Liu, J. Zeng, Y. Wang, S. A. Maitland, F. Idrizi, K. Ponnienselvan, L. J. Zhu, J. Luban, D. E. Bauer. Optimization of nuclear localization signal composition improves CRISPR-cas12a editing rates in human primary cells. GEN biotechnology, 1 (2022), pp. 271–284.

41. Y. Liu, R. S. Zou, S. He, Y. Nihongaki, X. Li, S. Razavi, B. Wu, T. Ha. Very fast CRISPR on demand. Science, 368 (2020), pp. 1265–1269, doi:10.1126/science.aay8204.

42. T. Zor, Z. Selinger. Linearization of the Bradford protein assay increases its sensitivity: theoretical and experimental studies. Analytical biochemistry, 236 (1996), pp. 302–308.

43. X. Yu, X. Liang, H. Xie, S. Kumar, N. Ravinder, J. Potter, X. de Mollerat du Jeu, J. D. Chesnut. Improved delivery of Cas9 protein/gRNA complexes using lipofectamine CRISPRMAX. Biotechnology letters, 38 (2016), pp. 919–929.

44. J. Schindelin, I. Arganda-Carreras, E. Frise, V. Kaynig, M. Longair, T. Pietzsch, S. Preibisch, C. Rueden, S. Saalfeld, B. Schmid. Fiji: an open-source platform for biological-image analysis. Nature methods, 9 (2012), pp. 676–682.

45. A. N. Shah, C. F. Davey, A. C. Whitebirch, A. C. Miller, C. B. Moens. Rapid reverse genetic screening using CRISPR in zebrafish. Nature methods, 12 (2015), pp. 535–540.

46. B. Li, N. Ren, L. Yang, J. Liu, Q. Huang. A qPCR method for genome editing efficiency determination and single-cell clone screening in human cells. Scientific reports, 9 (2019), pp. 1–15.

47. K. Clement, H. Rees, M. C. Canver, J. M. Gehrke, R. Farouni, J. Y. Hsu, M. A. Cole, D. R. Liu, J. K. Joung, D. E. Bauer. CRISPResso2 provides accurate and rapid genome editing sequence analysis. Nature biotechnology, 37 (2019), pp. 224–226.

48. P. Mali, L. Yang, K. M. Esvelt, J. Aach, M. Guell, J. E. DiCarlo, J. E. Norville, G. M. Church. RNA-guided human genome engineering via Cas9. Science, 339 (2013), pp. 823–826.

49. J. Salsman, G. Dellaire. Precision genome editing in the CRISPR era. Biochemistry and cell biology, 95 (2017), pp. 187–201.

50. I. Maggio, H. A. Zittersteijn, Q. Wang, J. Liu, J. M. Janssen, I. T. Ojeda, S. M. van der Maarel, A. C. Lankester, R. C. Hoeben, M. A. Gonçalves. Integrating gene delivery and gene-editing technologies by adenoviral vector transfer of optimized CRISPR-Cas9 components. Gene therapy, 27 (2020), pp. 209–225.

51. R. Torres-Ruiz, M. Martinez-Lage, M. C. Martin, A. Garcia, C. Bueno, J. Castaño, J. C. Ramirez, P. Menendez, J. C. Cigudosa, S. Rodriguez-Perales. Efficient recreation of t (11; 22) EWSR1-FLI1+ in human stem cells using CRISPR/Cas9. Stem cell reports, 8 (2017), pp. 1408–1420.

52. S. Niinuma, Y. Wake, Y. Nakagawa, T. Kaneko. Importance of nuclear localization signal-fused Cas9 in the production of genome-edited mice via embryo electroporation. Biochemical and Biophysical Research Communications, 685 (2023), pp. 149140.

53. M. A. Tyumentseva, A. I. Tyumentsev, V. G. Akimkin. Protocol for assessment of the efficiency of CRISPR/Cas RNP delivery to different types of target cells. PloS one, 16 (2021), pp. e0259812.

54. A. Kagita, M. S. Lung, H. Xu, Y. Kita, N. Sasakawa, T. Iguchi, M. Ono, X. H. Wang, P. Gee, A. Hotta. Efficient ssODN-mediated targeting by avoiding cellular inhibitory RNAs through precomplexed CRISPR-Cas9/sgRNA ribonucleoprotein. Stem Cell Reports, 16 (2021), pp. 985–996.

55. R. Jayavaradhan, D. M. Pillis, M. Goodman, F. Zhang, Y. Zhang, P. R. Andreassen, P. Malik. CRISPR-Cas9 fusion to dominant-negative 53BP1 enhances HDR and inhibits NHEJ specifically at Cas9 target sites. Nature communications, 10 (2019), pp. 2866.

56. D. Lainšček, V. Forstnerič, V. Mikolič, Š. Malenšek, P. Pečan, M. Benčina, M. Sever, H. Podgornik, R. Jerala. Coiled-coil heterodimer-based recruitment of an exonuclease to CRISPR/Cas for enhanced gene editing. Nature Communications, 13 (2022), pp. 3604.

57. C. Brees, M. Fransen. A cost-effective approach to microporate mammalian cells with the Neon Transfection System. Analytical biochemistry, 466 (2014), pp. 49–50.

58. B. van de Kooij, A. Kruswick, H. van Attikum, M. B. Yaffe. Multi-pathway DNA-repair reporters reveal competition between end-joining, single-strand annealing and homologous recombination at Cas9-induced DNA double-strand breaks. Nature Communications, 13 (2022), pp. 5295.

59. Y.-W. Fu, X.-Y. Dai, W.-T. Wang, Z.-X. Yang, J.-J. Zhao, J.-P. Zhang, W. Wen, F. Zhang, K. C. Oberg, L. Zhang. Dynamics and competition of CRISPR–Cas9 ribonucleoproteins and AAV donor-mediated NHEJ, MMEJ and HDR editing. Nucleic acids research, 49 (2021), pp. 969–985.

60. H. Yang, S. Ren, S. Yu, H. Pan, T. Li, S. Ge, J. Zhang, N. Xia. Methods favoring homology-directed repair choice in response to CRISPR/Cas9 induced-double strand breaks. International journal of molecular sciences, 21 (2020), pp. 6461.

61. N. Arnoult, A. Correia, J. Ma, A. Merlo, S. Garcia-Gomez, M. Maric, M. Tognetti, C. W. Benner, S. J. Boulton, A. Saghatelian. Regulation of DNA repair pathway choice in S and G2 phases by the NHEJ inhibitor CYREN. Nature, 549 (2017), pp. 548–552.

62. S. V. Vartak, S. C. Raghavan. Inhibition of nonhomologous end joining to increase the specificity of CRISPR/Cas9 genome editing. The FEBS journal, 282 (2015), pp. 4289–4294.

63. R. Gupta, K. Somyajit, T. Narita, E. Maskey, A. Stanlie, M. Kremer, D. Typas, M. Lammers, N. Mailand, A. Nussenzweig. DNA repair network analysis reveals shieldin as a key regulator of NHEJ and PARP inhibitor sensitivity. Cell, 173 (2018), pp. 972–988. e923.

64. G. Li, D. Liu, X. Zhang, R. Quan, C. Zhong, J. Mo, Y. Huang, H. Wang, X. Ruan, Z. Xu. Suppressing Ku70/Ku80 expression elevates homology-directed repair efficiency in primary fibroblasts. The international journal of biochemistry & cell biology, 99 (2018), pp. 154–160.

65. M. Liu, S. Rehman, X. Tang, K. Gu, Q. Fan, D. Chen, W. Ma. Methodologies for improving HDR efficiency. Frontiers in genetics, 9 (2019), pp. 691.

66. W. C. Skarnes, E. Pellegrino, J. A. McDonough. Improving homology-directed repair efficiency in human stem cells. Methods, 164 (2019), pp. 18–28.

67. G. Li, H. Wang, X. Zhang, Z. Wu, H. Yang. A Cas9–transcription factor fusion protein enhances homology-directed repair efficiency. Journal of Biological Chemistry, 296 (2021), pp.

68. J. Song, D. Yang, J. Xu, T. Zhu, Y. E. Chen, J. Zhang. RS-1 enhances CRISPR/Cas9-and TALEN-mediated knock-in efficiency. Nature communications, 7 (2016), pp. 10548.

69. W. Sun, H. Liu, W. Yin, J. Qiao, X. Zhao, Y. Liu. Strategies for enhancing the homology-directed repair efficiency of CRISPR-cas systems. The CRISPR journal, 5 (2022), pp. 7–18.

70. T. Kato-Inui, G. Takahashi, S. Hsu, Y. Miyaoka. Clustered regularly interspaced short palindromic repeats (CRISPR)/CRISPR-associated protein 9 with improved proof-reading enhances homology-directed repair. Nucleic Acids Research, 46 (2018), pp. 4677–4688.

71. K. Suzuki, Y. Tsunekawa, R. Hernandez-Benitez, J. Wu, J. Zhu, E. J. Kim, F. Hatanaka, M. Yamamoto, T. Araoka, Z. Li. In vivo genome editing via CRISPR/Cas9 mediated homology-independent targeted integration. Nature, 540 (2016), pp. 144–149.

