## Supplementary materials for "Optimizing Genome Editing in Bovine Cells: A Comparative Study of Cas9 Variants and CRISPR Delivery Methods"

### Supplementary Table S1. Sequences of oligos.

| Oligo name | Sequence (5’-3’) | Size (bp) |
|---|---|---|
| *Sry*-F | CTGCCAGGACGTATTGAGGG | 898 |
| *Sry*-R | AAAGAGCGCCTTTGTTAGCG | |
| *Sry*-NGS-F | TCGTCGGCAGCGTCAGATGTGTATAAGAGACAGACAGTGCAGTCGTATGCTTCT | 200 |
| *Sry*-NGS-R | GTCTCGTGGGCTCGGAGATGTGTATAAGAGACAGTTCATGGGTCGCTTGACGTG | |
| *Sry*-OUT-F | CTGCACAGCCTACGCATCTA | 148 |
| *Sry*-OUT-R | TTGGTACCCAAGCAGTTCCC | |
| *Sry*-ON-F | AACAGTGCAGTCGTATGCTT | 90 |
| *Sry*-ON-R | GCGAGAGTAGTTTGTTGCTGT | |
| *Sry* sgRNA1 | CGATGTTTACAGTCCAGCTG | |
| *Sry* sgRNA2 | GACCTCGTCGGAGAGCCAAG | |
| *Sry* donor ssODN | GCTATGTTCAGAGTATTGAACGACGATGTTTACAGTCCAGTAAGTGACTAGGTAACTGAGTAGCCTGTGGTACAGCAACAAACTACTCTCGCTTTTAGGAAAGA | |
| *Nanos2*-F | AATGGCTCTGGGGTGTATGG | 910 |
| *Nanos2*-R | CGAGTCCAGAGATGGGGAGA | |
| *Nanos2*-NGS-F | TCGTCGGCAGCGTCAGATGTGTATAAGAGACAGAAAAGGGGGTGCAGTTCCTC | 211 |
| *Nanos2*-NGS-R | GTCTCGTGGGCTCGGAGATGTGTATAAGAGACAGCCTTGGGTCTCCAACCCTTG | |
| *Nanos2*-OUT-F | CAGGAAAGACCCCCTTCCAC | 148 |
| *Nanos2*-OUT-R | CAGGAGCAGAGAGGAACTGC | |
| *Nanos2*-ON-F | CCATCAGCTGCTCCTGTCTG | 177 |
| *Nanos2*-ON-R | CCTTGGGTCTCCAACCCTTG | |
| *Nanos2* donor ssODN | ACCAACACTCCCCCGGGTGCCATGCAGCTGCCACCCTTTGTAAGTGACTAGGTAACTGAGTAGCACATGTGGAAGGACTACTTCAACCTGAGCCAAGTGGTGCT | |
| *Nanos2* sgRNA1 | GTAGTCCTTCCACATGTCAA | |
| *Nanos2* sgRNA2 | AGTCTCTCTACCGCCGCAGT | |
| *PRLR*-F | CCCTCCAAAGAACACACGGA | 859 |
| *PRLR*-R | AGTGCATAAAACCTGCGGGA | |
| *PRLR*-NGS-F | TCGTCGGCAGCGTCAGATGTGTATAAGAGACAGGGCCAATGGACCCAAATCTTC | 213 |
| *PRLR*-NGS-R | GTCTCGTGGGCTCGGAGATGTGTATAAGAGACAGCTCTGCTTGGTTGCCTTTCC | |
| *PRLR*-OUT-F | CCCTCCAAAGAACACACGGA | 126 |
| *PRLR*-OUT-R | ATGGGCCTGAGGTTCATCAC | |

| | | |
|---|---|---|
| *PRLR*-ON-F | ACCACAACATTGCTGACGTG | 141 |
| *PRLR*-ON-R | TCTGCTTGGTTGCCTTTCCT | |
| *PRLR*-sgRNA1 | GTCTGTTTGGTCCAGCGAAG | |
| *PRLR*-sgRNA2 | TGGATCTCCACGTATTCCAA | |
| *PRLR*-donor ssODN | TGTGAGCTGGCCCTGGGCATGGCCGGCACCACAGGCACTTCGCTGGACCAAACAGACCAACATGTTTAAGTGACTAGGTAACTGAGTAGCCGCTGGACCAAACAGACCAACATGCTTTAAAAGCCTCAAA | |
| Stop-codon-F | TAAGTGACTAGGTAACTGAGTAGC | |

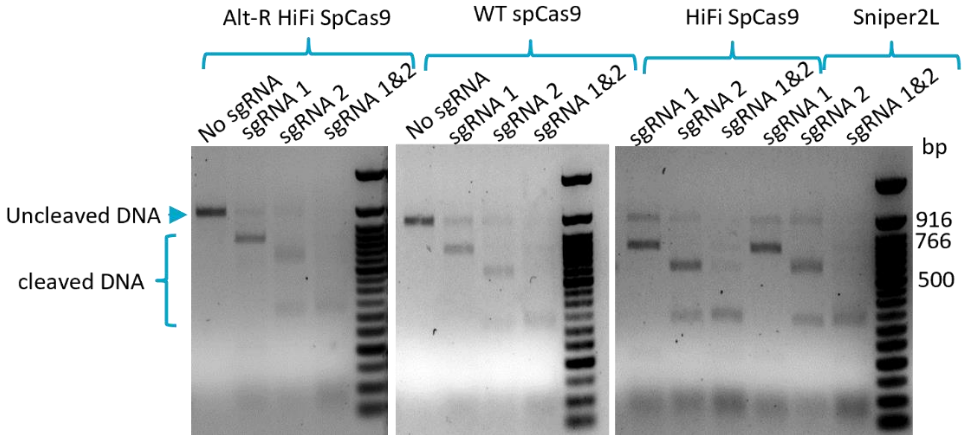


**Supplementary Figure S1. *In vitro* DNA cleavage assay with Cas9 nucleases.** Agarose gel electrophoresis showing the target DNA cleavage activity of the commercial Alt-R HiFi SpCas9, the in-house produced wild-type (WT) SpCas9, HiFi SpCas9 and Sniper2L. *Sry*-sgRNA1, *Sry*-sgRNA2, and target DNA amplified with primers *Sry*-F + *Sry*-R and genomic DNA extracted from WT MDBK cells were used for the assay.
